# Effects of anesthetic tricaine on stress and reproductive aspects of South American silver catfish (*Rhamdia quelen*) male

**DOI:** 10.1101/759340

**Authors:** Nathalia dos Santos Teixeira, Lis Santos Marques, Rômulo Batista Rodrigues, Darlan Gusso, Ana Amélia Nunes Fossati, Danilo Pedro Streit

## Abstract

Anesthesia is a common practice used in fish research and aquaculture. For both applications, it is important to understand anesthetics effects on the animal and tissues of interest to ensure the validity of data and to improve animal welfare. Captive fish production is only possible with artificial reproduction, and it is known that manipulation is a stressor stimulus in fish. The most common method of determining fish stress responses is measuring the circulating level of cortisol. Therefore, the purpose of this study was to evaluate the effects of different concentrations (100, 200, and 300 mg L^-1^) of the anesthetic tricaine methanesulfonate (MS-222) on cortisol levels and their influence on the sperm quality maintenance in *Rhamdia quelen.* After hormonal induction, 28 sexually mature males (average weight = 363.00 ± 71.24 g) were randomly distributed among treatments, and their semen and blood samples were collected. Anesthesia induction time, motility rate, sperm concentration and morphology, plasma cortisol levels, and reproductive hormones concentrations (testosterone, 17-α-hydroxyprogesterone, and estradiol) were evaluated. Anesthesia with 100 mg L^-1^ MS-222 presented a longer induction time than that with 200 and 300 mg L^-1^ MS-222. Sperm motility rate was significantly higher in the control than in the 300 mg L^-1^ treatment but did not differ among the control, 100, and 200 mg L^-1^ treatments. Estradiol level was significantly higher in non-anesthetized than in anesthetized fish, but plasma cortisol levels did not differ significantly between treatments (182.50 ± 42.03 ng mL^-1^). MS-222 anesthetizes fish by blocking the sodium channels, preventing the development of nerve action potentials. However, MS222 at concentrations of 100, 200, and 300 mg L^-1^ did not prevent stress in South American silver catfish males. In addition, its use did not maintain sperm quality, as it impaired motility and decreased levels of plasma estradiol.

## 1. Introduction

Stressors trigger a cascade of endocrine changes responsible for the stress response that can be divided into three main phases. The primary response is related to changes in the activity of neurotransmitters that lead to increased circulating levels of catecholamines and corticosteroids, mainly cortisol [1]. With the increase of these hormones in the circulatory system, several subsequent effects can be observed at blood and tissue levels, setting the secondary response. These changes include increase in circulating glucose and lactate levels, development of osmoregulatory disorders, and hematological and immunological changes [1, 2]. The persistence of these stressors, characterizing the tertiary response, may induce a significant impairment of well-being, negatively influencing physiological aspects such as immunity [3], behavior, and reproduction [2] in fish [4] and other species [5, 6, 7].

Animal reproduction is regulated by a complex interaction of various hormones that can be individually or collectively modulated by environmental and management factors [8, 9]. Thus, depending on the moment of the life cycle, severity, and stress duration, stress may affect reproduction [10]. The biological consequences of reproductive stress can be expressed by changes in both the reproductive behavior and the quantity and quality of the gametes [11], since they depend on an adequate hormonal environment during their development [12]. Thus, the importance of care in fish management should be taken into account, considering the health of the animals, the environmental sustainability of the aquaculture systems, and the profitability of the activity, since care is crucial to determine the survival success of the offspring in a production system.

Understanding the effects of stressful events at the population and individual level is indispensable for conservation biology, management of wild populations, and aquaculture [13]. Thus, one way to improve fish management is to minimize stress, preserving well-being. To achieve this goal, the use of anesthetics and/or sedatives has been suggested in fish farms [14].

Tricaine methanesulfonate (MS-222; ethyl 3-amino benzoate methanesulfonate), is currently the most commonly used drug in anesthesia, sedation, and euthanasia by immersion baths [15]. This is the only drug approved by the Food and Drug Administration (FDA) for use within fish for consumption in the US, although it is mandatory to wait at least 21 days for human consumption [2]. For fish anesthesia, the indicated concentration has a large variation among species that may vary from 50-400 mg L^-1^ of MS-222 [16].

To achieve satisfactory anesthesia and recovery in juveniles of South American silver catfish *Rhamdia quelen,* the indicated dose of MS-222 is 300 mg L^-1^ [17]. However, to date, no study has evaluated the effects of MS-222 anesthesia on seminal quality and reproductive hormonal profile of *R. quelen* males during reproductive management.

Thus, the objective of this study was to evaluate the effects of different MS-222 anesthetic concentrations (100, 200, and 300 mg L^-1^) on sexual steroids hormonal profile, sperm quality, and stress response (i.e., cortisol levels and differential leukocyte count) of *R. quelen* males during reproductive management.

## 2. Materials and methods

### 2.1. Fish maintenance and experimental conditions

Experimental protocols were performed according to Ethics and Animal Welfare Committee of the Federal University of Rio Grande do Sul (project number: 35840).

Two-year-old silver catfish (28 fish; 363.00 ± 71.24 g) were acclimated in seven plastic tanks (500 L) with a black background and a constant flow of aerated water for four weeks before the experiment, which was carried out during the summer. Fish were fed twice a day (8 h and 16 h) with a commercial diet (32 % crude protein, Acqua Fish, Supra^®^, Alisul, Brazil) until apparent satiety. Experimental parameters were as follows: water temperature, 27 ± 0.5 °C; pH, 6.8 ± 0.2; dissolved oxygen, 5.5 ± 0.5 mg L^-1^, and natural photoperiod.

### 2.2. Experimental design

The treatments consisted of increasing concentrations of MS-222 (Sigma-Aldrich, EUA, CAS Number: 886-86-2) for fish anesthesia (100, 200, and 300 mg L^-1^) and the control treatment where the animals were not anesthetized, including four treatments with seven replicates (each individual animal being considered as a repetition). The control group was exposed to anesthetic-free water and the duration time calculated from the mean of the other treatments.

### 2.3. Hormonal induction

Hormone induction was performed with intracavitary application of carp pituitary extract at the concentration of 3 mg kg^-1^ (pituitary / fish weight) using insulin syringe (1 mL) and a 13 × 0.45 mm needle. After 240 hours-degree, the fish were placed in tanks with 10 L of water containing the different anesthetic concentrations of each treatment.

### 2.4. Anesthetic baths

The anesthetic baths were performed individually for each fish prior to the collection of semen. Each fish was carefully removed from the maintenance tank and placed inside the vessel containing the anesthetic concentration to be tested. Anesthesia induction solutions were replaced with new ones after sedation of 14 animals ensuring that all fish were exposed to the correct concentrations of anesthetic. Anesthesia induction was observed according to the characteristics of fish anesthesia stages (Table 1) [18], with the objective of reaching the stage of deep anesthesia (IV), in which the animal loses the muscle tone as well as the balance, presenting a slow but regular opercular movement. The anesthesia induction time was monitored through a digital timer, quantifying the time each fish was immersed within the solution until the total loss of its movement and considerable decrease of the opercular movement.

**Table 1.**
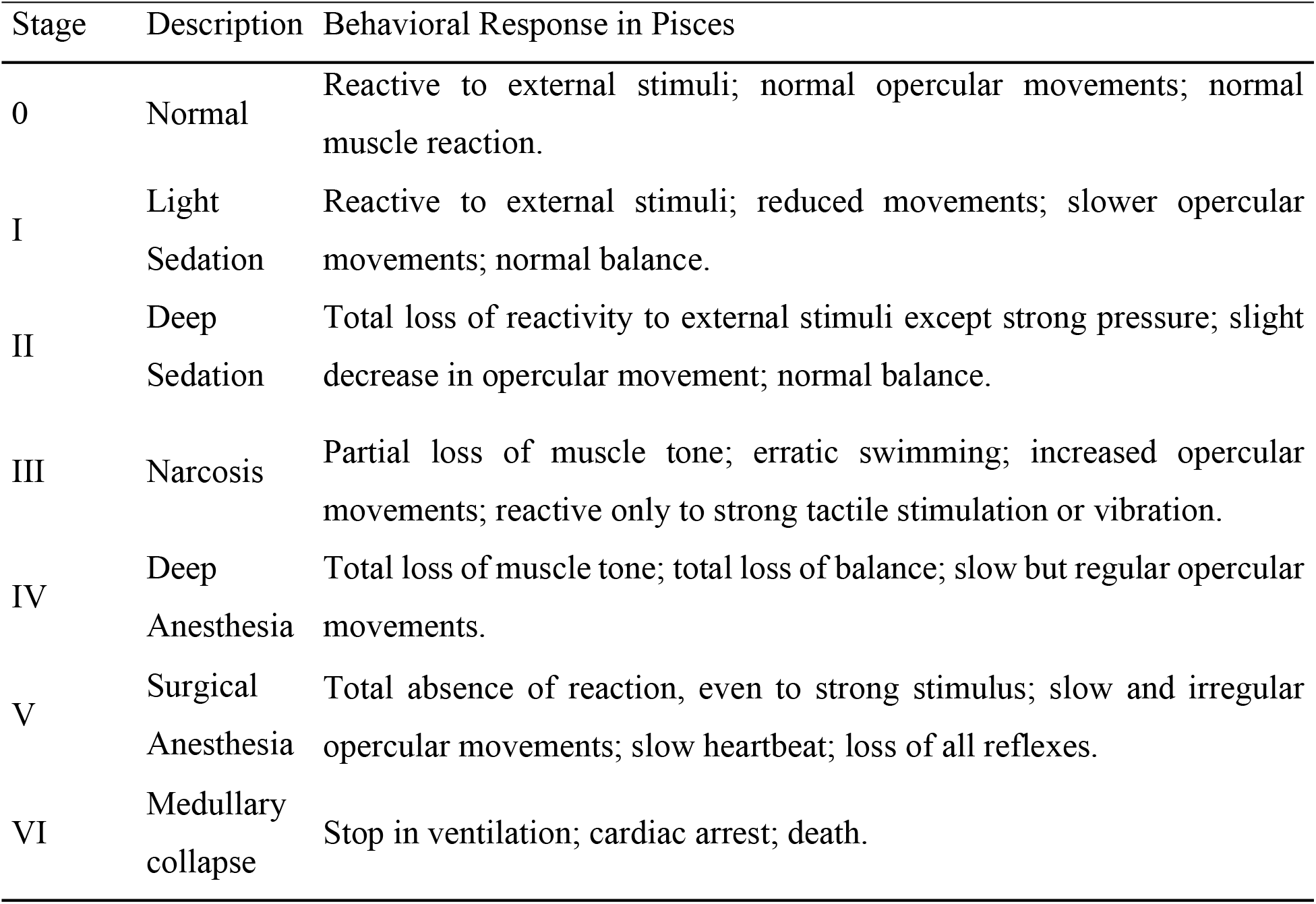
Stages of fish anesthesia

### 2.5. Semen quality collection and evaluation

For the collection of semen an anteroposterior massage was applied in the abdominal region with the fish slightly inclined with the head upwards, collecting the semen with the help of a 5 mL syringe. During collection, the first drop of semen was discarded and possible contamination with feces, blood, or urine was avoided.

#### 2.5.1. Membrane morphology, motility, and integrity analysis

For the evaluation of sperm morphology, samples were fixed in 10 % buffered formalin solution at 1:1000 dilution. The samples were subjected to the Rose Bengal staining (4 %) and analyzed under a light microscope (100×), with a total of 200 spermatozoa per smear. The percentage of normal and non-normal spermatozoa was measured, as well as the percentage of sperm abnormalities, i.e., primary pathologies (macrocephaly, microcephaly, head degenerations, fractured tail, strongly curled tail, simple bent tail) and secondary pathologies (free normal head, proximal droplet, distal droplet, coiled tail distal) [19].

For the evaluation of sperm motility, the semen was diluted and activated in 58 mM NaCl solution in a ratio of 2:20 μL, then a volume of 2 μL of this solution was placed on a histological slide for evaluation under an optical microscope (40×). Sperm motility rate was estimated in percentage. The percentage of cells with an intact membrane was evaluated using the eosin-nigrosin dyes. Semen was diluted (1:100) in Ginsburg Fish Ringer’s solution (sodium chloride – NaCl 6.50 g, potassium chloride – KCl 0.250 g, calcium chloride dihydrate – CaCl_2_ (H_2_O)_2_ NaHCO_3_ 0.20 g in 1000 mL of distilled water, pH 7.5, 300 mOsm) [20]. Subsequently, 20 μL of this dilution was mixed with 20 μL of the dye for a smear on histological slide and evaluation of 200 spermatozoa per slide. Sperm that had a non-stained head were considered intact and spermatozoa with a stained head were considered non-intact.

### 2.6. Collection and evaluation of blood parameters

#### 2.6.1. Blood collection

With the animal still anesthetized, the blood was collected with a 3 mL syringe and a 25 × 0.7 mm needle inserted into the ventral region, caudal to the genital region, at an angle of 45-90 ° towards the ventral region of the spinal cord to allow the puncture of the caudal vein. A maximum of 3 mL were collected from each animal.

#### 2.6.2. Measurement of hormone levels and leukocyte differential

Blood samples were transferred to microtubes (Microvette^®^ 500 μL, serum gel with clot activator, Sarstedt, Deutschland) for plasma separation. Cortisol levels and hormonal profile of sex steroids were determined by enzyme-linked immunosorbent assay (ELISA) according to the manufacturer’s instructions (17-β-estradiol, Testosterone, EIA Kit SYMBIOSIS DIAGNOSTIC LTDA, Brazil; 17-α-hydroxyprogesterone, Cortisol, EIA DBC Kit, Canada).

A blood smear was performed with 0.2 mL of each blood sample in order to determine the influence of the anesthetic on the defense cells [11]. The differential leukocyte count was performed according to literature [21].

### 2.7. Statistical analysis

The normality and homogeneity of the data were verified by the Shapiro-Wilk test and the Levene test, respectively. After verification of compliance with the statistical assumptions, data were analyzed by one-way ANOVA, followed by the Tukey averages comparison test, at 5 % significance level. Data that did not present normal distribution were analyzed by Kruskal-Wallis analysis, followed by Dunn’s test, at 5 % significance level. The data of variables presenting significant differences between treatments were subjected to regression analysis, and those that showed a significant coefficient were presented. Pearson’s correlation analysis was applied to verify the relationship between the variables analyzed. Statistical Analysis System 9.4 and GraphPad Prism 7.0 software was used to aid in statistical analysis and graphing.

## 3. Results

A significant difference *(p* = 0.0145) was observed for anesthesia induction time (s) among anesthetized fish with increasing concentrations of MS-222 (Fig 1). Animals anesthetized with 100 mg L^-1^ MS-222 presented the longest anesthesia induction time (440.14 ± 51.32 seconds), differing from fish anesthetized with 200 and 300 mg L^-1^, which presented anesthesia induction times of 283.43 ± 46.35 and 243, 38 ± 29.54 s, respectively.

**Fig 1.**
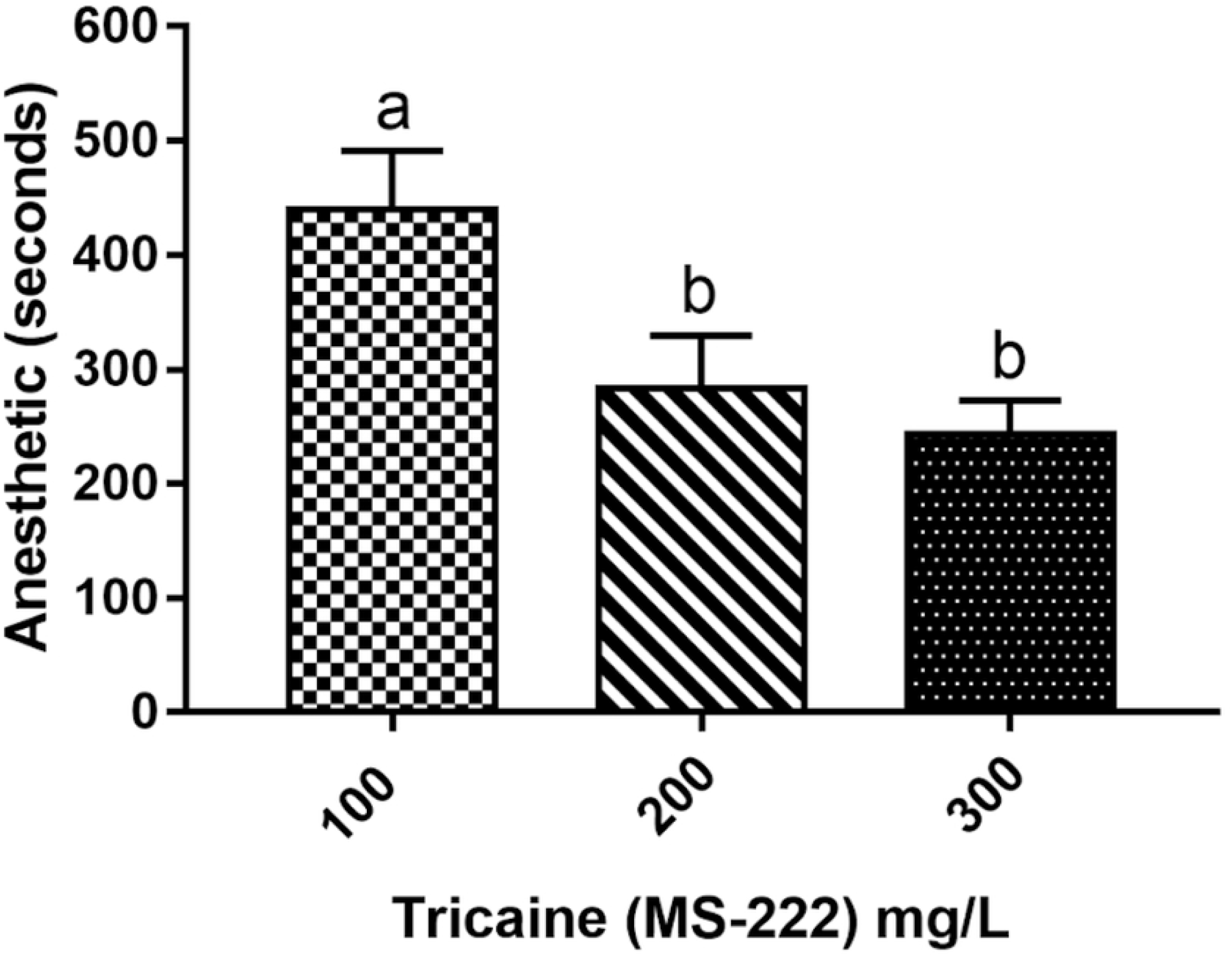
Anesthetic induction (seconds) of *R. quelen* anesthetized with increasing concentrations of tricaine (MS-222). Different letters indicate significant difference by the Tukey test (p<0.05). Data expressed as Mean±SEM.

Fish from the control group had the highest estradiol concentration (118.90 ± 21.22 ng mL^-1^), which was statistically different from that of those anesthetized with 100 (62.43 ± 10.92 ng mL^-1^), 200 (54.71 ± 7.55 ng mL^-1^) and 300 mg L^-1^ tricaine (61.75 ± 11.78 ng mL^-1^) (Fig 2A). No significant difference was observed among treatments for cortisol levels *(p* = 0.3394), testosterone *(p* = 0.3100) and 17-α-hydroxyprogesterone levels *(p* = 0.7176) (Fig 2B).

**Fig 2.**
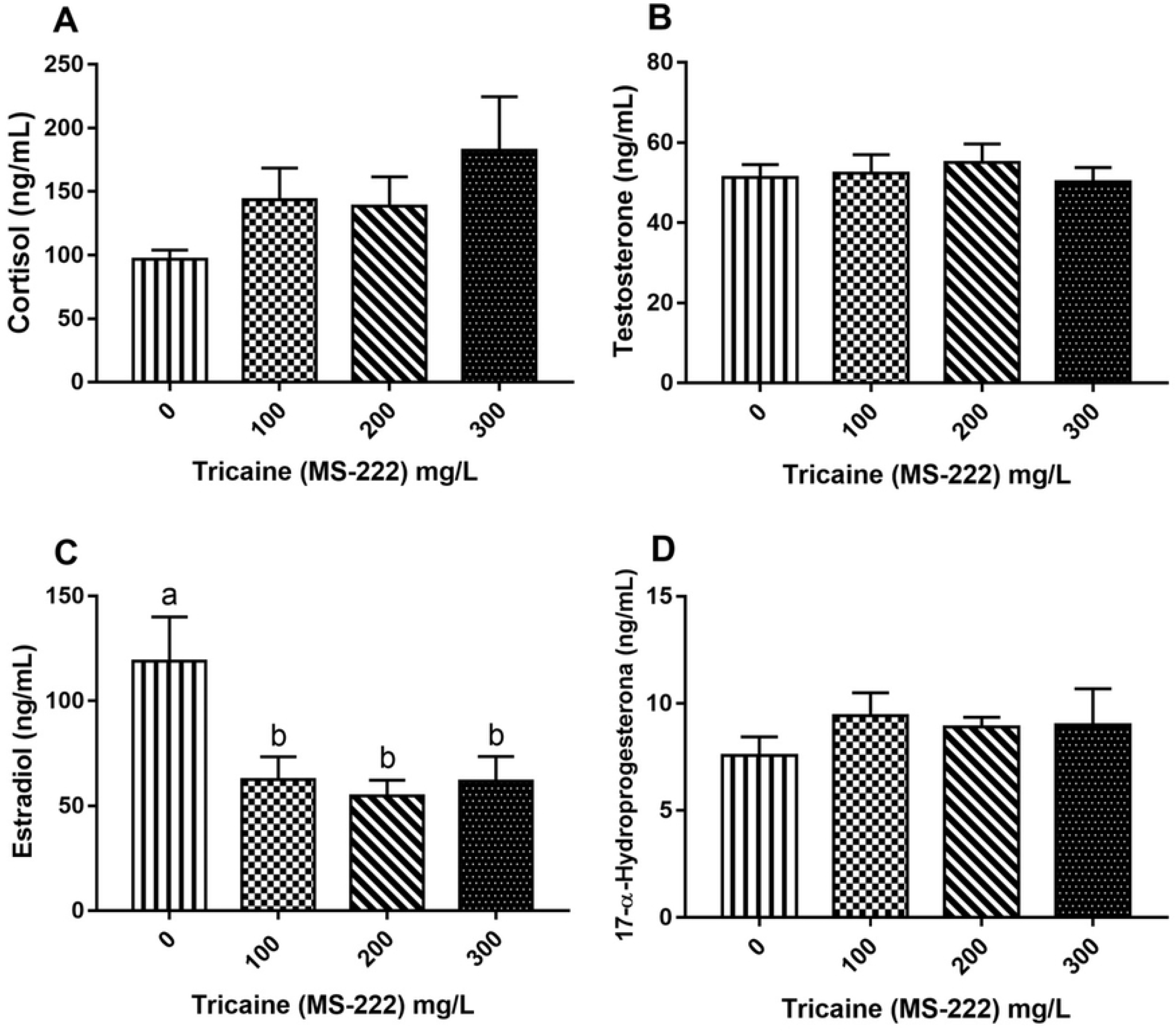
Hormone levels of *R. quelen* anesthetized with increasing concentrations of tricaine (MS-222). A: Cortisol (ng/mL); B: Testosterone (ng/mL); C: Estradiol (ng/mL); D: 17-α-Hydroxyprogesterone (ng/mL). Different letters indicate significant difference by the Tukey test *(p<0.05).* Data expressed as Mean±SEM.

No significant difference was observed among treatments in percent of lymphocytes *(p* = 0.5080), neutrophils *(p* = 0.2016), monocytes *(p* = 0.1527), and granular leukocytes *(p* = 0.4879) (Fig 3).

**Fig 3.**
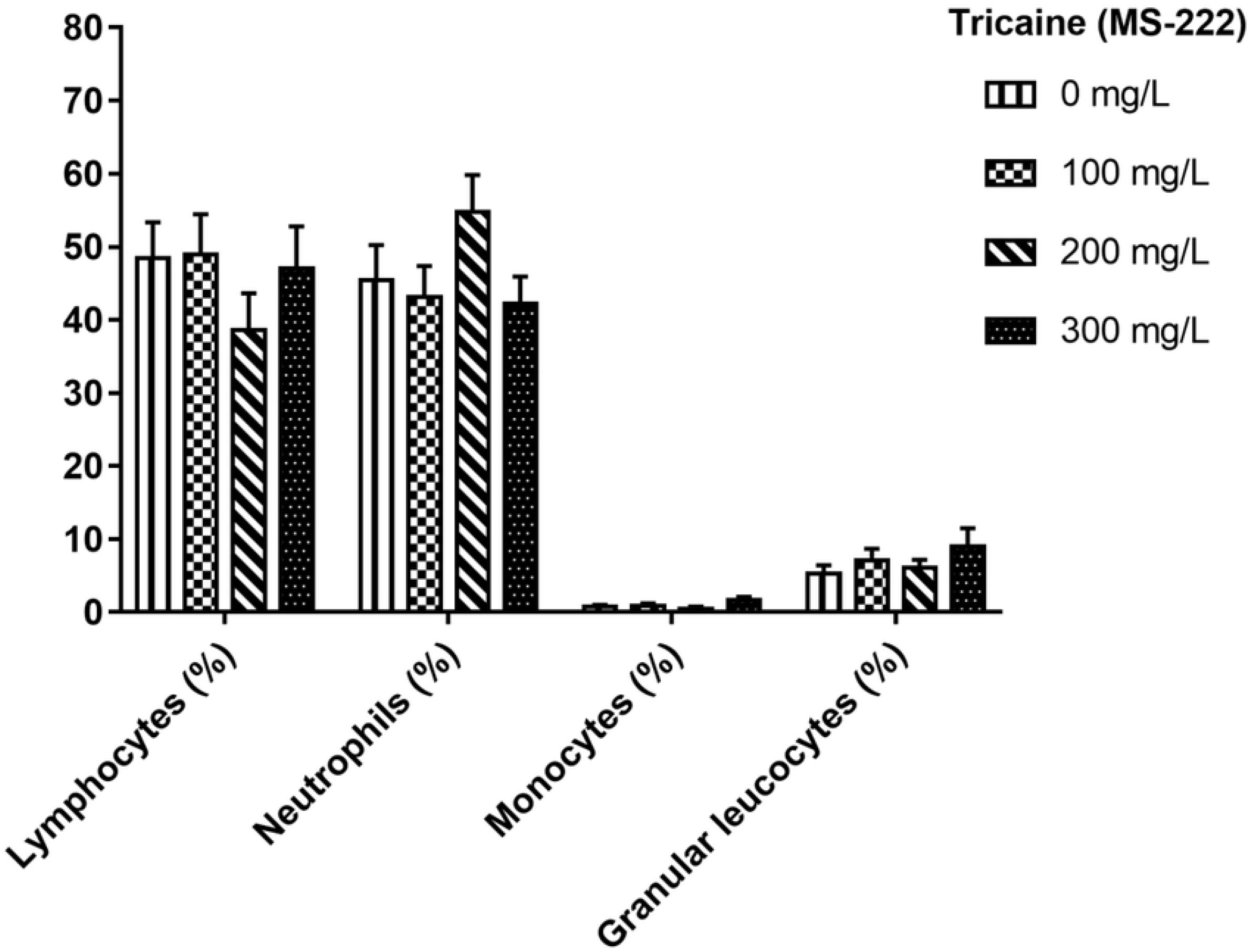
Leukogram of *R. quelen* anesthetized with increasing concentrations of tricaine (MS-222). There was no statistical difference among treatments (p>0.05). Data expressed as Mean±SEM.

A significant difference *(p* = 0.0324) was observed in sperm motility rate between animals anesthetized with the highest anesthetic concentration (300 mg L^-1^, 66.25 ± 5.6 %) and control animals (90.00 % ± 4.47 %). However, control animals did not differ statistically from the animals anesthetized with 100 and 200 mg L^-1^ MS-222 (80.00 % ± 4.36 % and 74.29 % ± 6.12 %, respectively) (Fig 4A). There was no difference among treatments in membrane integrity *(p* = 0.4135; Fig 4B) and morphology *(p* = 0.4430; Fig 4C).

**Fig 4.**
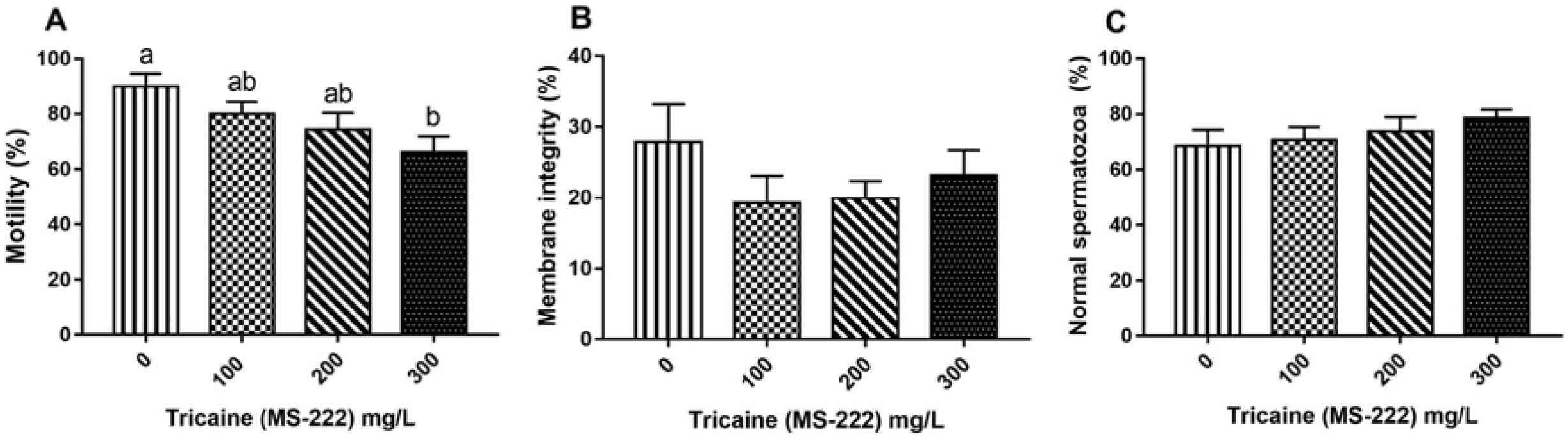
Qualitative variables of *R. quelen* semen anesthetized with increasing concentrations of tricaine (MS-222). A: Motility (%); B: Membrane integrity (%); C: Normal morphology (%). Different letters indicate significant difference by the Tukey test (p<0.05). Data expressed as Mean±SEM.

The variables of spermatic morphology are shown in Table 2. There was no significant difference among treatments, showing that anesthesia did not influence the percent of spermatozoa with simple bent tail, fractured tail, strongly coiled tail, coiled tail distal, proximal droplet, distal droplet, macrocephaly, microcephaly, head degeneration, and free normal head.

**Table 2.** Sperm morphology in *R. quelen* semen anesthetized with increasing concentrations of tricaine (MS-222).

| Variables (%) | Tricaine (MS-222) |  |  |  | Value of p |
| --- | --- | --- | --- | --- | --- |
|  | 0 mg/L | 100 mg/L | 200 mg/L | 300 mg/L |  |
| Simple bent tail | 8.29±1.52 | 8.57±1.87 | 7.79±2.19 | 5.69±1.05 | 0.5923* |
| Fractured tail | 1.93±1.06 | 2.00±0.53 | 1.71±0.34 | 1.81±0.54 | 0.7922** |
| Strongly coiled tail | 6.29±2.57 | 6.07±1.33 | 4.29±1.57 | 5.31±1.93 | 0.8840* |
| Coiled tail distal | 4.14±1.02 | 5.50±1.69 | 4.93±1.22 | 4.94±2.08 | 0.9498* |
| Proximal droplet | 0.21±0.15 | 0.21±0.15 | 0.36±0.24 | 0.44±0.29 | 0.9944** |
| Distal droplet | 4.14±2.57 | 4.07±1.90 | 3.43±1.43 | 0.63±0.37 | 0.2037** |
| Macrocephaly | 1.57±1.19 | 0.43±0.23 | 0.36±0.14 | 0.31±0.21 | 0.8551** |
| Microcephaly | 0.64±0.42 | 0.07±0.07 | 0.71±0.63 | 0.19±0.09 | 0.6253** |
| Head degeneration | 3.86±1.48 | 2.07±0.59 | 2.14±0.92 | 1.63±0.49 | 0.8073** |
| Free normal head | 0.29±0.10 | 0.93±0.43 | 0.57±0.35 | 0.44±0.20 | 0.7289** |

\* Not significant by analysis of variance ( $p>0.05$ ). \*\* Not significant by analysis Kruskal-Wallis ( $p>0.05$ ). Data expressed as Mean±SEM.

Regression models were applied on the variables that showed a significant difference among the treatments (Fig 5). Increasing concentrations of MS-222 showed a linear negative effect on anesthetic induction and motility rate, i.e., the higher the anesthetic concentration for *R. quelen*, the shorter the time fish took to enter the anesthesia induction stage and the lower the sperm motility rate. Estradiol presented a quadratic (polynomial) negative response to increasing levels of MS-222, i.e., as the anesthetic concentration for *R. quelen* increased, lower blood estradiol levels where maintained until reaching a point where estradiol levels began to rise again. According to the quadratic regression equation, the MS-222 concentration value where the hormone reached its minimum was 204.41 mg L^-1^.

**Fig 5.**
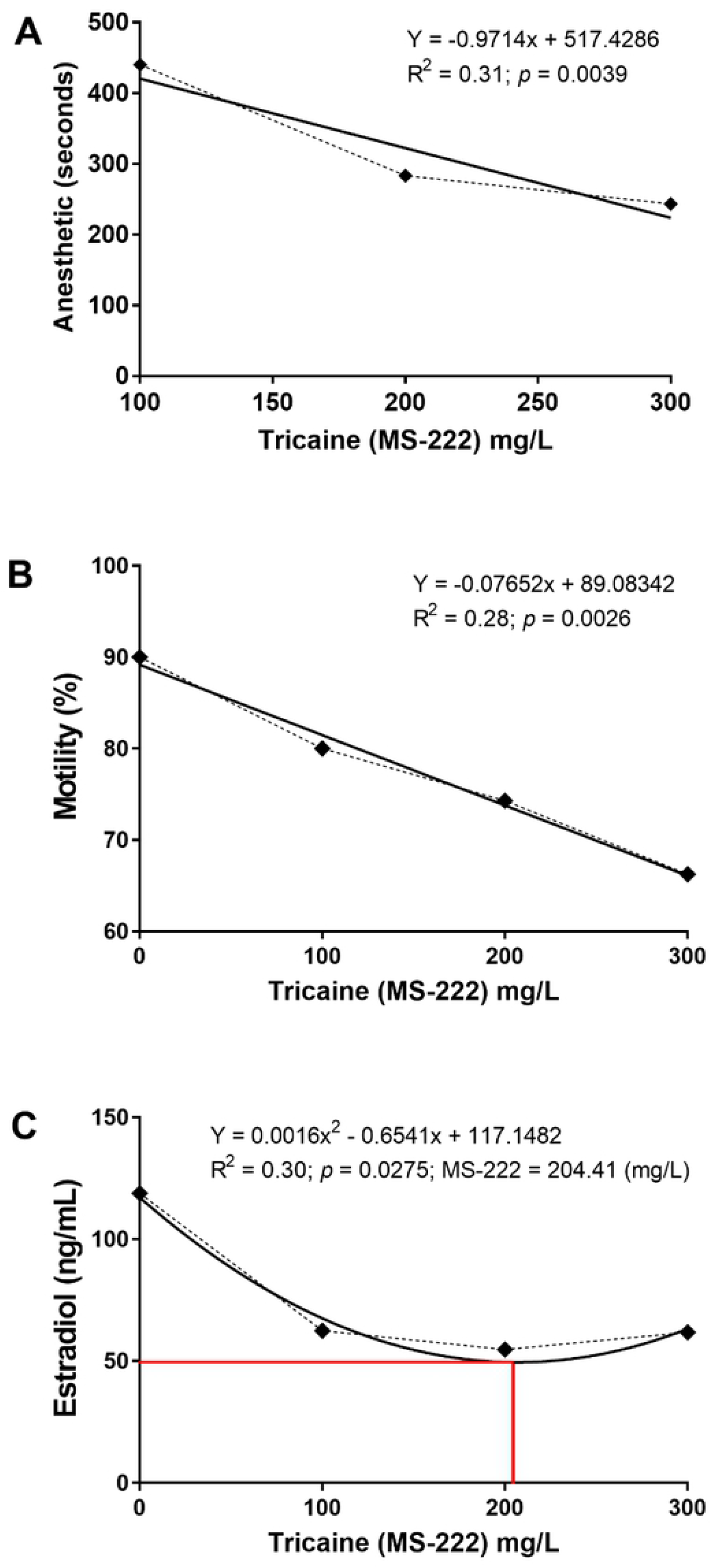
Graphical representation of the regression analysis that presented statistical significance. A: Induction of anesthesia (seconds); B: Motility rate (%); C: Estradiol (ng/mL).

Pearson’s correlation analysis was applied to all variables to detect variables that were related to each other, and graphs presenting the significant correlations are shown in Fig 6. In the correlation analyses, we observed a negative correlation between anesthesia induction time and percent of lymphocytes (R^2^ = −0.39, *p* < 0.05), percent of lymphocytes and neutrophils (R^2^ = −0.94, *p* < 0.001), percent of lymphocytes and monocytes (R^2^ = −0.44, *p* < 0.05), and between the percent of lymphocytes and granular leukocytes (R^2^ = −0.45, *p* < 0.05). A positive correlation was observed between the percent of neutrophils and anesthesia induction time (R^2^ = 0.46, *p* < 0.05), and between estradiol and percent of monocytes (R^2^ = 0.56, *p* < 0.01).

**Fig 6.**
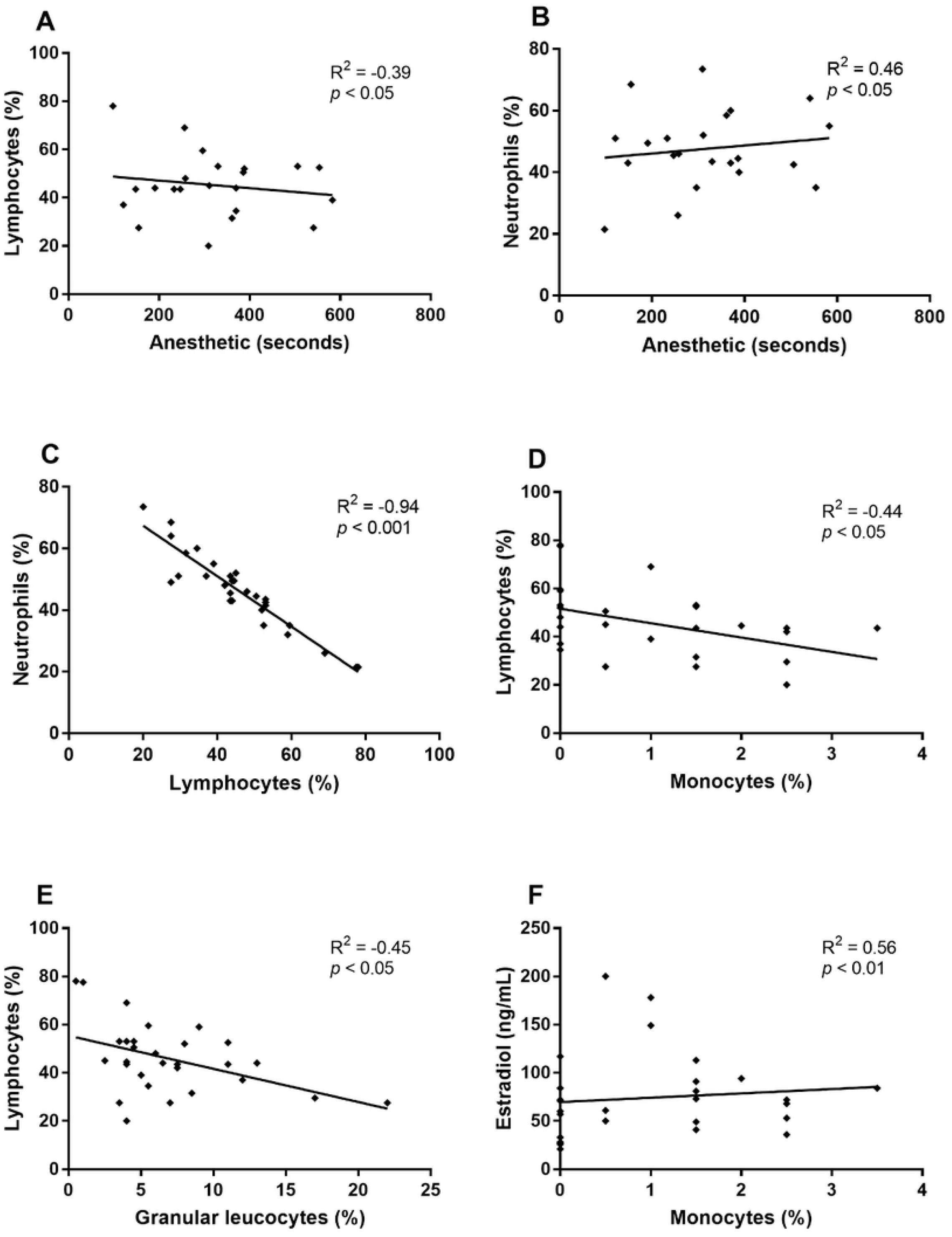
Graphical representation of Pearson correlation analyzes that presented statistical significance. A: Lymphocytes (%) x Induction of anesthesia (seconds); B: Neutrophils (%) x Induction of anesthesia (seconds); C: Neutrophils (%) x Lymphocytes (%); D: Lymphocytes (%) x Monocytes (%); E: Lymphocytes (%) x Granular leukocytes (%); F: Monocytes (%) x Estradiol (ng / mL).

## 4. Discussion

In this study, we evaluated the effects of different anesthetic (MS-222) concentrations on the sexual steroids hormonal profile, sperm quality maintenance, and stress response of *R. quelen* males during the reproductive management.

Anesthesia induction time for *R. quelen* decreased linearly with MS-222 concentration increase. Biological factors such as life cycle stages, age, size, weight, lipid content, and health need to be considered during the use of anesthetics, since the different biological factors are strictly correlated [22] and could cause different reactions and increased fish susceptibility to anesthetics. It is known that resistance and tolerance to different anesthetics vary among individuals and species [23]. In the present study, an increase in cortisol values was observed accompanying the increase in anesthetic concentration. Although no significant difference was detected, this alteration may be related to the possible irritation caused by the anesthetic bath. Moreover, studies have shown that the use of MS-222 (100 mg L^-1^) for juvenile *Ictalurus punctatus* increased cortisol over the basal concentration after 5 min of sedation [24]. In this study, the mean time of anesthesia using 100 mg L^-1^ considerably exceeded 5 min, which may have contributed to the blood cortisol increase in these fish. In addition, during exposure to MS-222, as the anesthetic begins to take effect, loss of balance may cause a stress response [15].

In both human and veterinary medicine, a sedative is often administered prior to anesthesia for the purpose of calming the patient and reducing the stress that the anesthetic or anesthetic procedure may cause [15].

The basal cortisol concentration in *R. quelen* males is 15.86 ng/mL, after acute stress, the concentration reaches 158.12 ng/mL [25] and the peak values range from 158.0 (males) to 207.0 ng/mL (females), 1 h after handling the animals [11]. In the present study, animals in the control group had a mean cortisol value of 96.86 ± 7.08 ng/mL, identifying a state of acute stress that may have been caused by hormonal induction approximately 10 h prior to collection of semen and blood. In addition, the concentrations of MS-222 tested did not reduce plasma cortisol concentration, consistent with the results of a previous study [26], where MS-222 was not able to prevent the increase of cortisol in *Pimephales promelas.* It is possible that an increased response to stress during the anesthetic procedure may have occurred due to the low availability of oxygen caused by insufficient gill ventilation or by direct stimulation of the hypothalamic–pituitary–interrenal axis (HHI) [27].

Currently, there is still insufficient knowledge to define when the limits of homeostatic fluctuations have been exceeded, leading to the state of stress [10]. Thus, although the circulating cortisol level is the most evaluated indicator in order to measure the stress response in fish, it should be interpreted in conjunction with other physiological variables [28]. Therefore, since the amplitude of physiological parameters in fishes has not yet been fully elucidated and it is important to consider as many variables as possible for determining the presence of stress, cortisol analysis alone is not the best way of evaluating the response to stress.

It is believed that the deleterious effects of stress on the immune response are preferentially mediated by the suppressive effects of glucocorticoids (i.e. cortisol) and are a consequence of the inability to adapt to chronic stressors [29]. Contrastingly, fish immune response against an acute stress is to enhance innate function to prepare the immune system for challenges such as the neuroendocrine systems for fight-or flight [30].

After evaluating *R. quelen* hematological parameters, it was identified that thrombocytes and lymphocytes are the most frequent organic defense cells, lymphocytes being the largest representative of leukocytes [31]. Previous research [32] has demonstrated that acute stress may cause a reduction in the number of circulating lymphocytes, monocytes and special granulocytic cells (SGC) in *R. quelen,* as well as the potential increase in percentage of circulating neutrophils. In the present study, no significant difference was observed among treatments for differential leukocyte count.

The administration of exogenous cortisol results in lower oocyte growth, condition factor, and plasma levels of testosterone and 17-β-estradiol in *Oreochromis mossambicus* [33]. A study carried out with *R. quelen* females showed that the high level of plasma cortisol resulted in lower concentrations of plasma 17-β-estradiol [8]. The authors have suggested the possibility of an inhibitory effect of estradiol on aromatase, an enzyme that converts testosterone to estradiol. This fact corroborates the results obtained in the present study for the determination of plasma 17-β-estradiol, found in a lower concentration in the animals with a higher level of plasma cortisol [8].

The increase in cortisol may be associated with sperm motility reduction, since it was observed that in males of *Oncorhynchus mykiss* that cortisol level increase in fish confined alone in small tanks was related to a 10 % reduction in sperm motility [34]. It is also possible that stress hormones may interfere with plasma osmolarity and impair sperm quality [12]. Acute stress during capture and transport resulted in dilution of blood plasma and semen osmolality in mature males *Morone chrysops*, consequently decreasing motility rate [35]. Osmolarity is an important aspect of sperm quality, since it is a determinant factor for activation of sperm motility [36, 37]. In addition to cortisol, anesthetics may also interfere with this parameter, since direct contact between MS-222 and sperm decreased the sperm motility time of *O. mykiss* [28]. In the present study, sperm motility presented a negative linear response, suggesting that stress-related hormones have induced osmoregulatory dysfunctions [38] causing blood plasma dilution and alteration in seminal plasma osmolarity [37], causing pre-activation of spermatozoa and, consequently, decreased sperm motility [35].

The use of MS-222 for anesthesia of *R. quelen* breeding herds influenced many of the evaluated variables, and several variables correlated with each other as a response to MS-222. We observed a negative correlation between anesthesia induction time and the percentage of lymphocytes, as well as a positive correlation between anesthesia induction time and the percentage of neutrophils. The results show that leukocytes are sensitive to the changes caused by stressful factors of reproductive management and the anesthetic procedure. Thus, hormone levels and differential leukocyte counts could be good indicators of the efficacy of MS-222 anesthesia in male *R. quelen* breeding. However, for a better understanding of MS-222 influence on the reproduction of *R. quelen* males, we suggest that further studies test other anesthetic concentration ranges and evaluate other reproductive and stress response parameters.

At the concentrations studied (100, 200, and 300 mg L^-1^), the anesthetic MS-222 was not efficient for decreasing plasma cortisol levels. In addition, there was a decrease in sperm motility rate and a reduction in plasma levels of estradiol. Therefore, anesthesia with MS-222 at these concentrations is not indicated for *R. quelen* breeding males.

## Acknowledgements

This research was supported by Conselho Nacional de Desenvolvimento Científico e Tecnológico (CNPq).

